# Neuroinflammation links the neurogenic and neurodegenerative phenotypes of *Nrmt1^-/-^* mice

**DOI:** 10.1101/2025.09.02.673860

**Authors:** James P. Catlin, Shane Fraher, Jessy J. Alexander, Christine E. Schaner Tooley

## Abstract

It is widely thought that age-related damage is the single biggest contributing factor to neurodegenerative diseases. However, recent studies are beginning to indicate that many of these diseases may have developmental origins that become unmasked overtime. It has been difficult to prove these developmental origins, as there are still few known links between defective embryonic neurogenesis and progressive neurodegeneration. We have created a constitutive knockout mouse for the N-terminal methyltransferase NRMT1 (*Nrmt1*^-/-^ mice). *Nrmt1*^-/-^ mice display phenotypes associated with premature aging. Specifically in the brain, they exhibit age-related striatal and hippocampal degeneration, which is accompanied by impaired short and long-term memory. These phenotypes are preceded by depletion of the postnatal neural stem cell (NSC) pools, which appears to be driven by their premature differentiation and migration.

However, this differentiation is often incomplete, as many resulting neurons cannot permanently exit the cell cycle and ultimately undergo apoptosis. Here, we show that the onset of apoptosis corresponds to increased cleavage of p35 into the CDK5 activator p25, which can promote neuroinflammation. Accordingly, *Nrmt1*^-/-^ brains exhibit an increase in pro-inflammatory cytokine signaling, astrogliosis, complement activation, microgliosis, and markers of a compromised blood brain barrier, all of which indicate an activated neuroimmune response. We also find *Nrmt1*^-/-^ mice do not activate a corresponding anti-inflammatory response. These data indicate that abnormal neurogenesis can trigger neuroinflammation, which in the absence of compensatory anti-inflammatory signaling, could lead to neuronal apoptosis and progressive neurodegeneration.

## Background

While it has long been thought that damage during the aging process is one of the main drivers of progressive neurodegenerative diseases (1), recent data suggests these diseases could result from developmental defects that become unmasked over time (2, 3). It is now generally accepted that Huntington’s disease (HD) has a developmental component (4), as Juvenile HD affects neurodevelopment, expression of Huntingtin CAG repeats in human embryonic stem cells (hESCs) results in misregulation of germ layer patterning, and increasing CAG repeat length alters neural differentiation (2, 5–7). Though not yet as widely accepted as HD, data are also accumulating that Parkinson’s disease (PD) and Alzheimer’s disease (AD) may have developmental origins as well. Many proteins known to be involved in PD (alpha-synuclein, VPS35, PINK1, LRRK2) and AD (ApoE, tau, APP) progression also have known neurogenic roles (8–14). In addition, there are many common signaling pathways that play key roles in both neurogenesis and neurodegeneration, including TGFβ, Wnt, and Notch signaling (15–17).

In order for neurodegenerative diseases to arise from early neurogenic defects, there must be compensatory mechanisms in place that can mask symptoms until later in life. It has been proposed that these mechanisms could include re-routing neuronal circuitry to circumvent neurons with weakened connectivity, utilization of partially redundant signaling pathways, and the interplay between pro-and anti-inflammatory responses (18–20). In the case of AD, the balance between pro-and anti-inflammatory signaling has been proposed to delay the onset of symptom presentation (21). Early in AD, anti-inflammatory M2 microglia can clear amyloid beta (Aβ) peptides through APOE/TREM2/ITAM/SYK-mediated phagocytosis (21–24). However, as Aβ levels accumulate, they can promote TLR4/NF-κB/complement signaling and the activation of pro-inflammatory M1 microglia, which leads to synapse engulfment and neuronal loss (21, 25–27). In this way, M2 activation by low Aβ load, could prevent neuronal death and delay pathological onset until the Aβ load is insurmountable and M1 activation predominates.

A better understanding of these potential compensatory mechanisms has been hindered by a lack of appropriate model systems that exhibit both neurogenic defects and progressive neurodegeneration. Though recent data from N-terminal RCC1 methyltransferase 1 (NRMT1) knockout mice (*Nrmt1^-/-^*), suggest these mice could help address this question. *Nrmt1^-/-^* mice exhibit developmental defects, including reduced size and a low survival rate past six-months-old (28). However, those that survive past six months experience phenotypes associated with premature aging, including early graying, loss of hair, kyphosis, and liver degeneration (28).

They also exhibit neurodegenerative phenotypes, including impaired short-and long-term memory and decreased striatal and hippocampal volume at three-months-old (29). These two regions are unique in that they contain, or are adjacent to, the two major neural stem cell (NSC) niches. Accordingly, at postnatal day 14 (P14), *Nrmt1^-/-^* mice exhibit a depletion of both NSC pools and a corresponding increase in intermediate progenitor and neuroblast populations (29). The neuroblasts appear to differentiate and migrate correctly, though they fail to exit the cell cycle and ultimately begin undergoing apoptosis by six weeks (29).

This combination of early alteration in NSC differentiation and progressive neurodegeneration positions *Nrmt1^-/-^* mice as a potential new model system for studying the link between these two processes. As these mice have a fairly rapid onset of neurodegeneration, they will also be useful for studying lack of potential compensatory mechanisms. Here, we assay whether the inability of neurons to permanently exit the cell cycle in *Nrmt1^-/-^* mice triggers either a pro-or anti-inflammatory response before the onset of neurodegeneration begins. We show that corresponding to the onset of apoptosis at six weeks, *Nrmt1^-/-^* mice also exhibit increased cleavage of p35 into the CDK5 activator p25 and increased levels of phosphorylated tau, a substrate of the CDK5/p25 complex. This coincides with an increase in pro-inflammatory cytokine signaling, astrogliosis, complement activation, microgliosis, and markers of a compromised blood brain barrier, which all indicate an activated pro-inflammatory response. In comparison, we see an inability to increase expression of the anti-inflammatory markers IL-10, IL-4, and TGFβ, indicating neurodegeneration may proceed quickly in these mice due to the inability to mount an anti-inflammatory response. These studies demonstrate that neuroinflammation can link abnormal neurogenesis and neurodegeneration, which may proceed more quickly in the absence of compensatory anti-inflammatory signaling.

## Material and Methods

### Maintenance of *Nrmt1^-/-^* Mice

Homozygous C57BL/6J-*Nrmt1*^-/-^ mice were bred and genotyped as previously described (28). Both female and male mice were used in these studies as no striking differences between male and female mice were observed. Euthanasia was performed by inhalation of carbon dioxide in a clear container followed by cervical dislocation or by perfusion with 4% paraformaldehyde (PFA) following anesthesia. Both the breeding and euthanasia protocols are approved by the SUNY Buffalo Animal Care and Use Committee.

#### Immunohistochemistry

Mice were anesthetized with isoflurane and perfused with 1X cold PBS, followed by 4% PFA. Brains were collected and post-fixed in 4% PFA for 72 hours and cryo-protected in 30% sucrose at 4°C for 1 week. 30 μm coronal sections were generated using a Microm HM 505 N cryostat (Thermo Fischer Scientific, Waltham, MA) and frozen onto Superfrost Plus microscope slides (Fisher Scientific). Samples were washed with 1x PBS and permeabilized/blocked in 1X PBS-T (2% Triton-X) containing 5% Normal Horse Serum for 1 hour at room temperature (RT).

Samples were incubated overnight at 4°C in 1x PBS containing 2% normal horse serum and primary antibodies at the following dilutions: rabbit anti-glial fibrillary acidic protein (GFAP, 1:500; Sigma-Aldrich, G9269), rabbit anti-ionized calcium-binding adaptor molecule 1 (Iba1, 1:100; Cell Signaling Technologies, 17198T), or FITC conjugated anti-complement C3 (1:200; Cappel, 55500). The following day, samples were incubated in 1x PBS containing 2% normal horse serum and secondary antibodies at the following dilutions: goat anti-rabbit Alexa Fluor 594 (1:500; Invitrogen, A11012) or goat anti-rabbit Alexa Fluor 647 (1:500; Invitrogen, A21245). Samples were then counterstained with Hoechst 33342 (1:2000; AnaSpec Inc., 83218) for 1 hour at RT. Samples were mounted with Fluoromount-G (Invitrogen, 00-4958-02). Images were captured with a Cytation-5 Multi-Mode reader (Biotek, Winooski, VT) or Yokogawa CSU-X1 spinning-disk Leica DMi600B microscope (Intelligent Imaging Innovations, Tokyo, Japan). For confocal microscopy, maximum Z-projections were generated using FIJI software (NIH).

Mean fluorescent intensity of whole image was measured using FIJI software, background intensity was subtracted from image and a manual threshold was set. The manual threshold was the same for each time-point and antibody pairing. Puncta counting was performed with ImageJ plug-in Puncta Process (GitHub) following background intensity subtraction. 10 regions of interest were generated (20 μm x 10 μm, 2000 μm total area analyzed) and centered around fluorescent puncta, find maxima threshold was set at 50.

### Immunoblotting

Brains were isolated from P14 and 6-week wild type (WT) and *Nrmt1^-/-^* mice and serial sections were generated using a Zivic Brain Matrix (Zivic Instruments, Pittsburgh, PA). The lateral ventricles and adjacent striatum were dissected and processed in lysis buffer containing Tris (pH 7.4, 50 mM), sodium chloride (150mM), NP-40 (1%), sodium deoxycholate (0.25%), EDTA (1mM), SDS (0.1%), sodium orthovanadate (6 mM), sodium fluoride (6 mM), beta-glycerophosphate (20 mM) and the protease inhibitors phenylmethylsulfonyl fluoride, aprotinin and leupeptin (MilliporeSigma). Lysates were centrifuged at 14,000 rpm for 25 min at 4°C. 10 μg of lysates were used for western blot analysis. Protein was separated on 12% SDS-polyacrylamide gels and transferred onto polyvinylidene difluoride membrane using the Trans-Blot Turbo Transfer System (Bio-Rad). Membranes were blocked in 5% milk solution made in Tris buffered saline containing 1% Tween (TBS-T) for 1 hour at RT. 5% BSA solution in TBS-T was used for phospho-specific anti-bodies. Membranes were blotted overnight at 4°C in 5% milk or 5% BSA solution containing primary antibodies at the following dilutions: rabbit anti-GFAP (1:1000; Sigma-Aldrich, G9269), rabbit anti-glyceraldehyde-3-phosphate dehydrogenase (GAPDH, 1:3000; Tocris, 2275-PC-100), rabbit anti-p35/p25 (1:1000; Cell Signaling Technologies), rabbit anti-phospho Tau (S202) (1:1000; Abcam, ab108387), or rabbit anti-β-tubulin (1:1000; Cell Signaling Technologies, 2128L). Blots were incubated in 5% milk or 5% BSA solution containing HRP-conjugated donkey anti-rabbit at 1:5000 (Jackson ImmunoResearch, 711-035-152), developed using Clarity Western ECL Substrate (Bio-Rad, 170-5060), and imaged on an ChemiDoc Imaging System (Bio-Rad). Protein quantification was performed with ImageJ software (NIH), with GAPDH or β-tubulin serving as loading controls.

### Quantitative Reverse Transcriptase PCR

Lateral ventricles and adjacent striatum were isolated from P14 and 6-week WT and *Nrmt1^-/-^* mice as previously described(29). Total RNA was extracted with TRIzol reagent (Thermo Fisher Scientific). cDNA was generated from 1μg of RNA using the SuperScript III first strand synthesis system (Thermo Fisher Scientific). iQ SYBR Green Supermix (Bio-Rad, 1708882) containing the following primers was used: TNFα Fwd 5’ - TCACCCACACCGTCAGCCGATTT - 3’, Rev 5’ - CACCCATTCCCTTCACAGAGCAA - 3’; IL-1β Fwd 5’ – GAAATGCCACCTTTTGACAGTG – 3’, Rev 5’ – TGGATGCTCTCATCAGGACAG – 3’; IL-6 Fwd 5’ – CTGCAAGAGACTTCCATCCAG – 3’, Rev 5’ - AGTGGTATAGACAGGTCTGTTGG – 3’; C1q Fwd 5’ – ATAAAGGGGGAGAAAGGGCT - 3’, Rev 5’ – CGTTGCGTGGCTCATAGTT - 3’; C1INH fwd 5’ – TTGAGTGCCAAGTGGAAGATAAC – 3’, Rev 5’ – GTGCTTTGGGAACACGGGTAC – 3’; C4 Fwd 5’ – ACCCCCTAAATAACCTGG – 3’, Rev 5’ – CCTCATGTATCCTTTTTGGA – 3’; Factor B Fwd 5’ – CTCCTCTGGAGGTGTGAGCG – 3’, Rev 5’ – GGTCGTGGGCAGCGTATTG – 3’; C3 Fwd 5’ – AGCAGGTCATCAAGTCAGGC – 3’, Rev 5’ – GATGTAGCTGGTGTTGGGCT – 3’; Factor H Fwd 5’ – CGTGAATGTGGTGCAGATGGG – 3’, Rev 5’ – AGAATTTCCACACATCGTGGCT – 3’; TGFβ; Fwd 5’-TACCATGCCAACTTCTGTCTGGGA-3’, Rev 5’ - ATGTTGGACAACTGCTCCACCTTG - 3’; IL-4 5’ – CTCGAATGTACCAGGAGCCA – 3’, Rev 5’ – CACATCCATCTCCGTGCATG – 3’; IL-10 Fwd 5’ – ATTTGAATTCCCTGGGTGAGAAG – 3’, Rev 5’ – CACAGGGGAGAAATCGATGACA – 3’; TIMP1 Fwd 5’ – GCATCTGGCATCCTCTTGTT – 3’, Rev 5’ – TGGGGAACCCATGAATTTAG – 3’; MMP9 Fwd 5’ - CATTCGCGTGGATAAGGAGT - 3’ Rev 5’ - ACCTGGTTCACCTCATGGTC – 3’; Claudin-5 Fwd 5’ – TTTCTTCTATGCGCAGTTGG – 3’, Rev 5’ – GCAGTTTGGTGCCTACTTCA – 3’; Hdgf Fwd 5’ – CATGAGAGCCTGTAGCCAC – 3’, Rev 5’ – GTGGGCTTAGAGGAGAGAG – 3’. Hepatoma derived growth factor (Hdgf) was used as the housekeeping gene. Fold change was determined with the ΔΔ CT quantification method.

## Results

### *Nrmt1^-/-^* mice exhibit upregulation of p25

The inability of neurons in the striatum of *Nrmt1^-/-^* mice to permanently exit the cell cycle is reminiscent of the neurons that re-enter the cell cycle during the early stages of AD (30).

Reinitiating the cell cycle in neurons causes them to exhibit progressive axon initial segment (AIS) loss, reduced dendritic spine density, and altered synaptic function (30). It has been hypothesized that this altered synaptic function results from hyperactivation of cyclin-dependent kinase 5 (CDK5) and a shift back to its cell cycle substrates at the cost of its targets at pre-and postsynaptic terminals of mature neurons, including Kv1 potassium channel subunits (30, 31).

To determine if *Nrmt1^-/-^* mice exhibit hyperactivated CDK5, we used Western blots to assay levels of its activators p35 and p25. In response to neurotoxic stress, p35 is cleaved into p25, which hyperactivates CDK5 and promotes cell death (32). Accordingly, high levels of the CDK5/p25 complex have been associated with the pathology of many neurodegenerative diseases, including AD, HD, and PD (33–36).

We found that at day P14, when over 10% of NeuN-positive cells in the striatum are actively in the cell cycle in *Nrmt1^-/-^* mice (29), p25 levels are significantly decreased as compared to wild type (WT) mice (**Fig. 1a**). In addition to its activation in pathological conditions, p25 is also involved in early oligodendrocyte differentiation (37, 38), so its low levels at P14 in *Nrmt1^-/-^* mice may indicate additional lineage specification defects. However, by 6 weeks, p25 levels have significantly increased in *Nrmt1^-/-^*mice and are almost undetectable in WT mice (**Fig. 1b**), indicating the presence of a neurotoxic stimuli specific to *Nrmt1^-/-^* mice. We also found that levels of tau protein phosphorylated at Ser 202, a known target of CDK5/p25 (39), significantly increase between P14 and 6 weeks (**Fig. 1c,d**), further verifying progressive CDK5 hyperactivation in *Nrmt1^-/-^* mice.

**Fig. 1.**
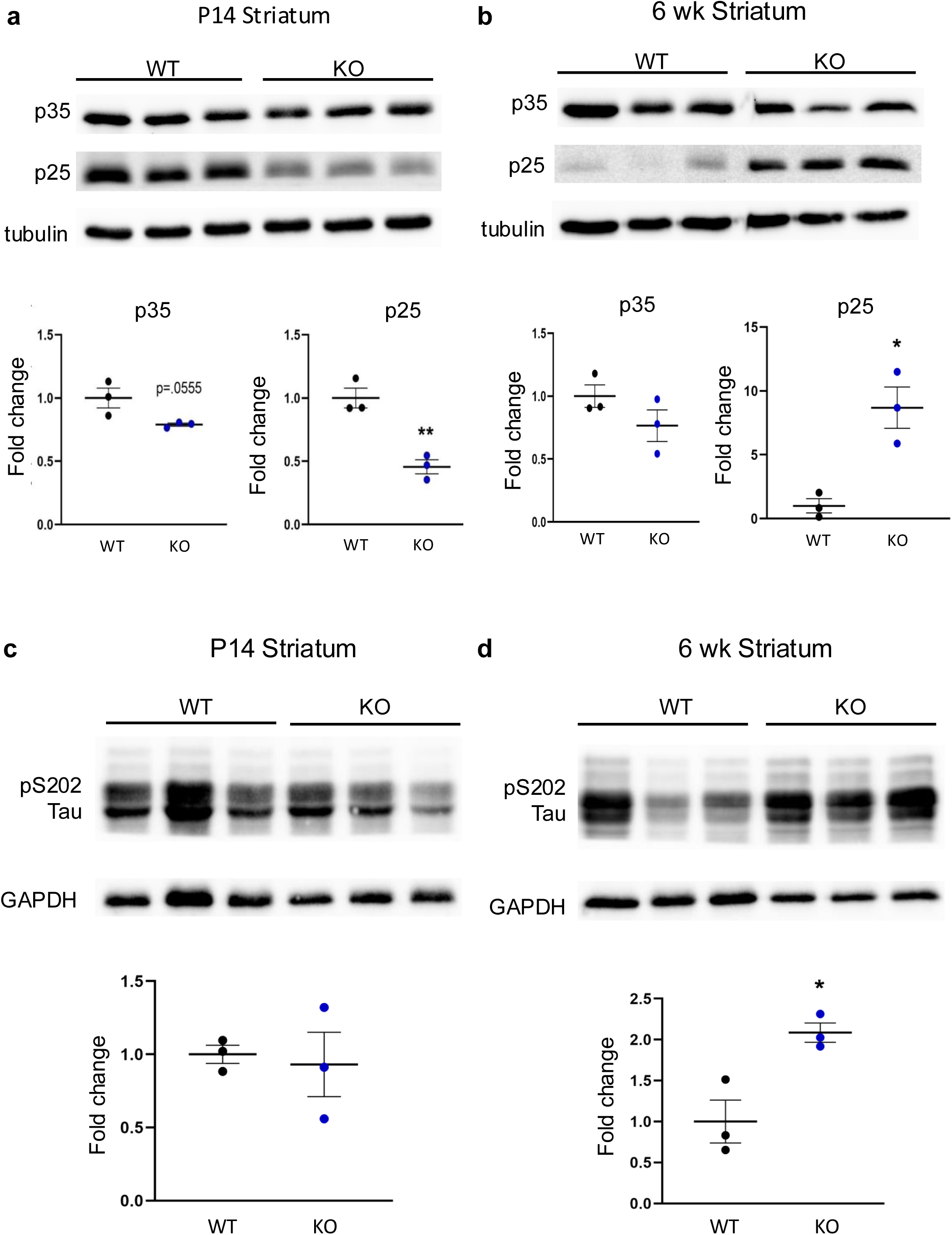
CDK5 hyperactivation in *Nrmt1^-/-^* mice. Western blots showing p25 protein expression is significantly decreased in **a**) P14 *Nrmt1^-/-^* mice but **b**) significantly increased at 6 weeks. Protein expression normalized to β-tubulin. Phosphorylation of the p25/CDK5 substrate Tau at serine 202 (pS202 Tau) is **c**) not altered in *Nrmt1^-/-^* mice at P14 but **d**) significantly increased at 6 weeks. Protein expression was normalized to GAPDH. *p<0.05, **p<0.01 as determined by unpaired t-test and error bars signify mean±SEM. n=3.

### Pro-inflammatory cytokine signaling and astrogliosis

One downstream consequence of increased p25 production is the initiation of a pro-inflammatory cytokine signaling cascade, including elevated expression of tumor necrosis factor-alpha (TNF-α), interleukin 1 (IL-1), and interleukin 6 (IL-6) (40, 41). To determine if *Nrmt1^-/-^*mice exhibit increased mRNA expression of these signaling molecules, we performed quantitative real-time PCR (qPCR) analysis on brain lysates isolated from P14 and 6-week *Nrmt1^-/-^* and WT mice. We found that TNFα expression was increased in *Nrmt1^-/-^* mice at both P14 and 6 weeks (**Fig. 2a**), which corresponds to TNFα being an initiating factor of the cytokine signaling pathway (42).

**Fig. 2.**
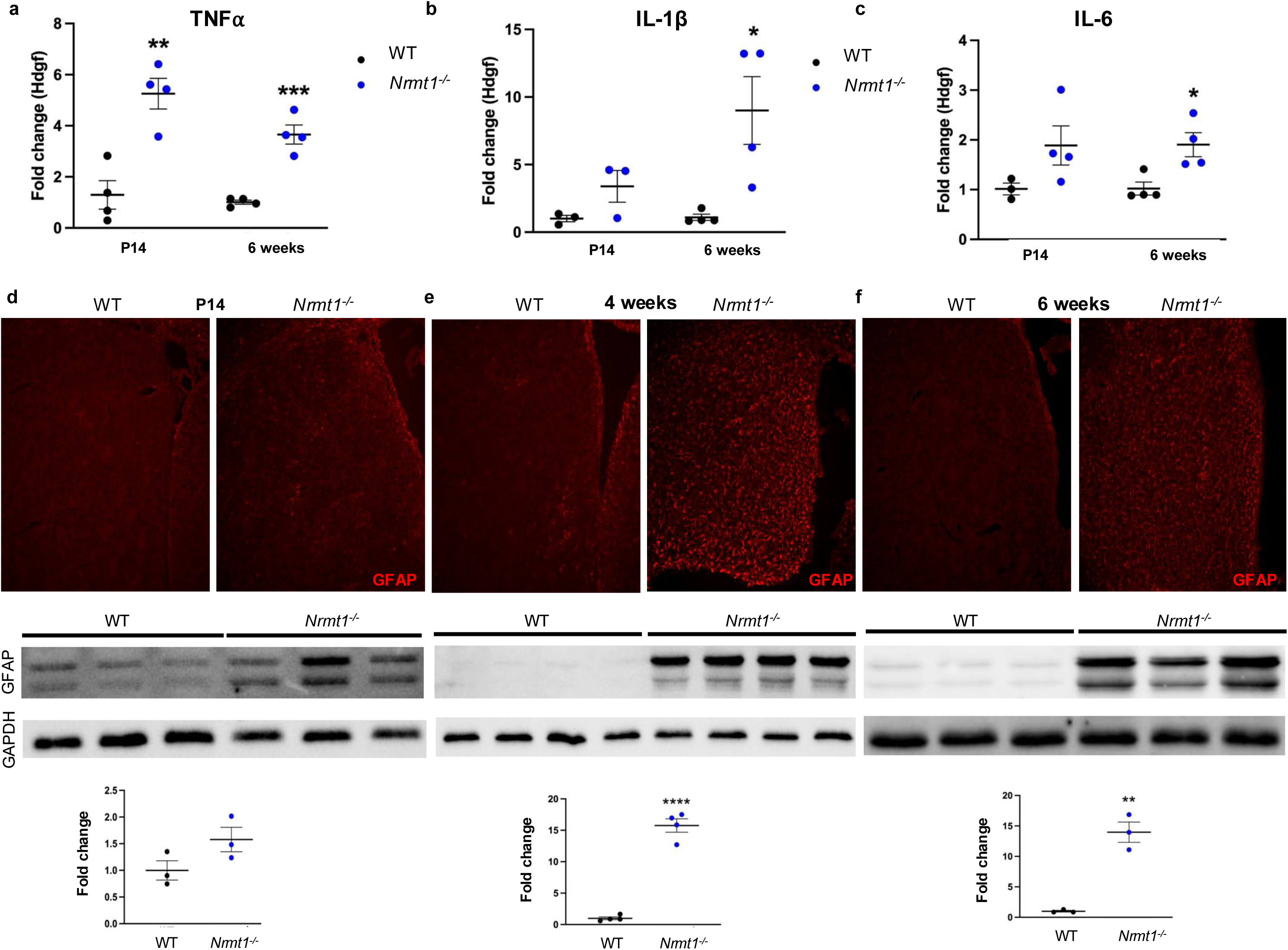
Cytokine signaling and astrogliosis. qPCR analysis showing that **a**) TNF⍺ expression is significantly increased in *Nrmt1^-/-^* mice at both P14 and 6 weeks, while **b,c**) IL-1β and IL-6 expression only significantly increases by 6 weeks. Immunohistochemistry and Western blots showing GFAP protein is **d**) not significantly increased in the striatum of *Nrmt1^-/-^* mice as compared to WT controls at P14. However, by **e**) 4 and **f**) 6 weeks, GFAP protein levels are significantly increased, indicating the initiation of astrogliosis. *p<0.05, **p<0.01, ****p<0.0001 as determined by unpaired t-test and error bars signify mean±SEM. n=3-4.

However, both IL-1β and IL-6, which are downstream of TNFα, do not significantly increase expression in *Nrmt1^-/-^* mice until 6 weeks (**Fig. 2b,c**). These data indicate that pro-inflammatory cytokine signaling begins in *Nrmt1^-/-^* mice by P14 with the activation of TNFα, and full-signaling through IL-1β and IL-6 is occurring by 6 weeks.

Another downstream consequence of increased p25 production and pro-inflammatory cytokine signaling is the activation of astrocytes (astrogliosis) (43). Astrocytes respond to neuronal activity to provide nutrients and recycle neurotransmitters, as well as mediate neuroinflammatory responses in the event of neuronal damage or dysfunction (44–46). To determine if p25 overexpression was promoting astrogliosis in *Nrmt1^-/-^*mice, confocal microscopy was used to assess glial fibrillary acidic protein (GFAP) immunoreactivity, a marker for activated astrocytes, within the striatum of *Nrmt1^-/-^* mice and WT controls. No difference in GFAP immunoreactivity in the striatum was observed between *Nrmt1^-/-^* and control mice at P14 (**Fig. 2d**). However, beginning at 4 weeks, there was an increase in GFAP immunoreactivity in the striatum of *Nrmt1^-/-^* mice (**Fig. 2e**), which persisted at 6 weeks (**Fig. 2f**). This is also when *Nrmt1^-/-^* mice first exhibit loss of striatal neurons (29). To confirm, striatal protein lysates from *Nrmt1^-/-^* and WT mice were analyzed via Western blot, and we observed a significant increase in GFAP protein expression at 4 weeks (**Fig. 2e**) and 6 weeks (**Fig. 2f**). These data indicate the presence of astrogliosis within the striatum of *Nrmt1^-/-^*mice.

### Complement activation and microgliosis

Astrogliosis can be both neuroprotective and neurotoxic, though coincidental expression of complement protein C3 indicates neurotoxicity (45). C3 is primarily produced by astrocytes, and its deposition along weakened synapses targets these synapses for removal through microglia-mediated phagocytosis (27, 45, 47, 48). C3 upregulation has been observed in post mortem human brains of neurodegenerative diseases (49, 50), and it has been proposed that complement-directed engulfment of synapses may be responsible for symptoms of early cognitive decline (47–52). To determine if the astrogliosis seen in *Nrmt1^-/-^* mice corresponds with increased C3 expression, we stained striatal sections with an antibody against C3. While we observed a significant increase in C3 expression in *Nrmt1^-/-^* mice at P14, 4 weeks, and 6 weeks, the staining pattern differed between ages (**Fig. 3**). At P14, C3 staining was diffuse (**Fig. 3a**) and co-stained with astrocyte processes (**Sup. Fig. 1**), reminiscent of the C3 staining of reactive astrocytes seen in traumatic brain injury (53). At 4 and 6 weeks, we observe punctate C3 immunoreactivity (**Fig. 3b,c**), indicative of synaptic deposition.

**Fig. 3.**
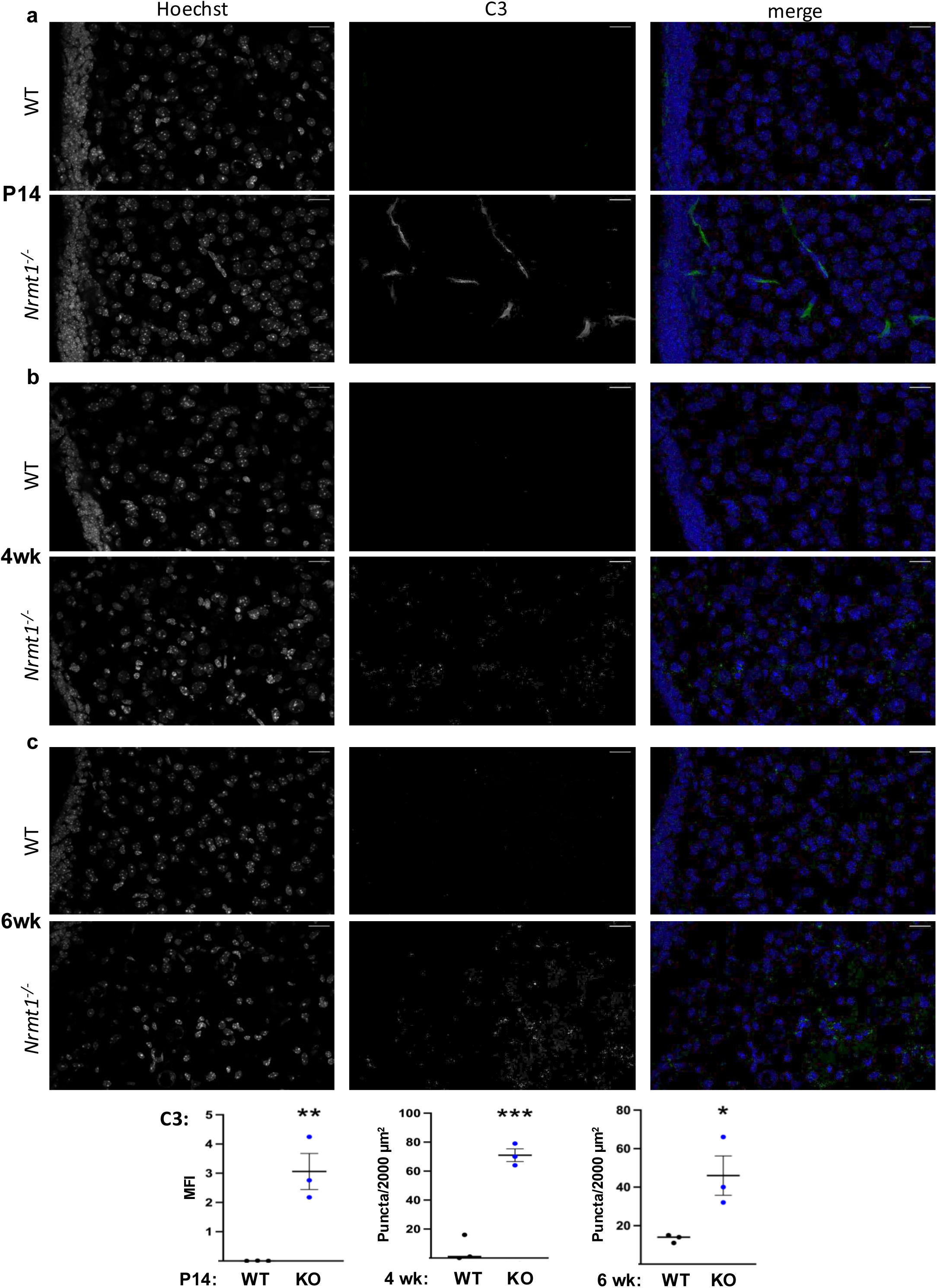
Complement C3 expression in the brain of *Nrmt1^-/-^* mice. **a** At two weeks C3 expression in *Nrmt1^-/-^* mice is significantly increased over wild type (WT), though exhibits a diffuse staining pattern. At **b**) 4 weeks and **c**) 6 weeks, C3 expression remains significantly increased, though the staining pattern is more punctate. Scale bars represent 20 μm unless otherwise stated. *p<0.05, **p<0.01, and ***p<0.001 as determined by unpaired t-test and error bars signify mean±SEM. n=3.

Microglia are the resident phagocytic cells of the brain and are responsible for clearing pathogens, dying cells, or weakened synapses after C3 deposition (47, 51, 54). To determine if the punctate C3 deposition correlates with microglia activation (microgliosis) in *Nrmt1^-/-^*mice, we co-stained striatal sections from 6-week *Nrmt1^-/-^* and WT mice with antibodies against C3 and ionized calcium-binding adaptor molecule 1 (IBA1), a marker of activated microglia (55). Using confocal microscopy, we observed a significant increase in IBA1 immunoreactivity within the striatum of *Nrmt1^-/-^* mice compared to WT controls (**Fig. 4a-f**) and found that many of the C3 puncta are within IBA1+ cells, suggesting that the microglia are undergoing active phagocytosis (**Fig. 4g**).

**Fig. 4.**
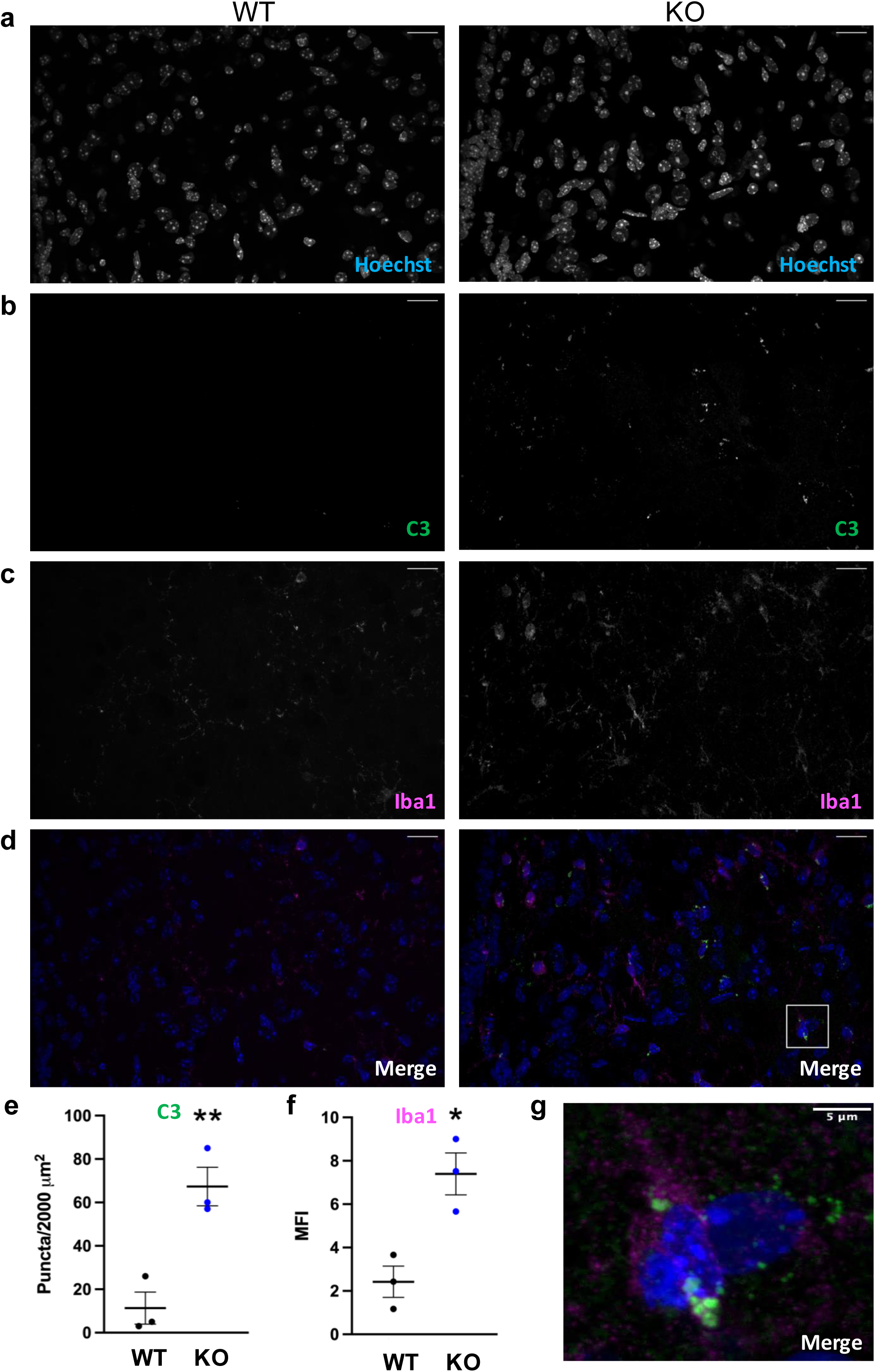
Co-localization of C3 and IBA1 staining. **a-f** Significantly increased C3 staining in the striatum of *Nrmt1^-/-^* mice at 6 weeks corresponds to a significant increase in IBA1 immunoreactivity. **g)** Inset showing an IBA1 positive cell that seems to have internalized some C3 puncta. Scale bars represent 20 μm unless otherwise stated. *p<0.05, **p<0.01 as determined by unpaired t-test and error bars signify mean±SEM. n=3.

To get a more comprehensive view of Complement activation, we performed qPCR to determine mRNA expression levels of Complement pathway members, including C1q, C1INH1, C3, C4, factor B (FB), and factor H (FH) in striatal lysate of P14 and 6-week WT and *Nrmt1^-/-^*mice. We observed upregulation of C1q, C1INH, and C4 as early as P14 (**Fig. 5a-c)**, implicating activation of the classical complement pathway. Components of the alternative pathway (C3, FB, FH), which amplify the classical signal, remained unaltered in striatal lysates of *Nrmt1^-/-^* mice at P14 (**Fig. 5d-f)**. However, mRNA expression of all three genes was significantly upregulated at 6-weeks (**Fig. 5d-f),** indicating complement signaling is increasing over time. Overall, components of the classical pathway are transcriptionally upregulated by P14, which coincides with the appearance of mitotic neurons (29). C3 puncta deposition is visible by 4 weeks (**Fig. 3b)**, which corresponds with early astrogliosis (**Fig. 2e),** and activation of the alternative pathway is seen by 6-weeks, when we also observe active microgliosis (**Fig. 4c,f**) and the onset of neuronal apoptosis (29). These data suggest that complement activity could be marking mitotic neurons for removal before overt loss.

**Fig. 5.**
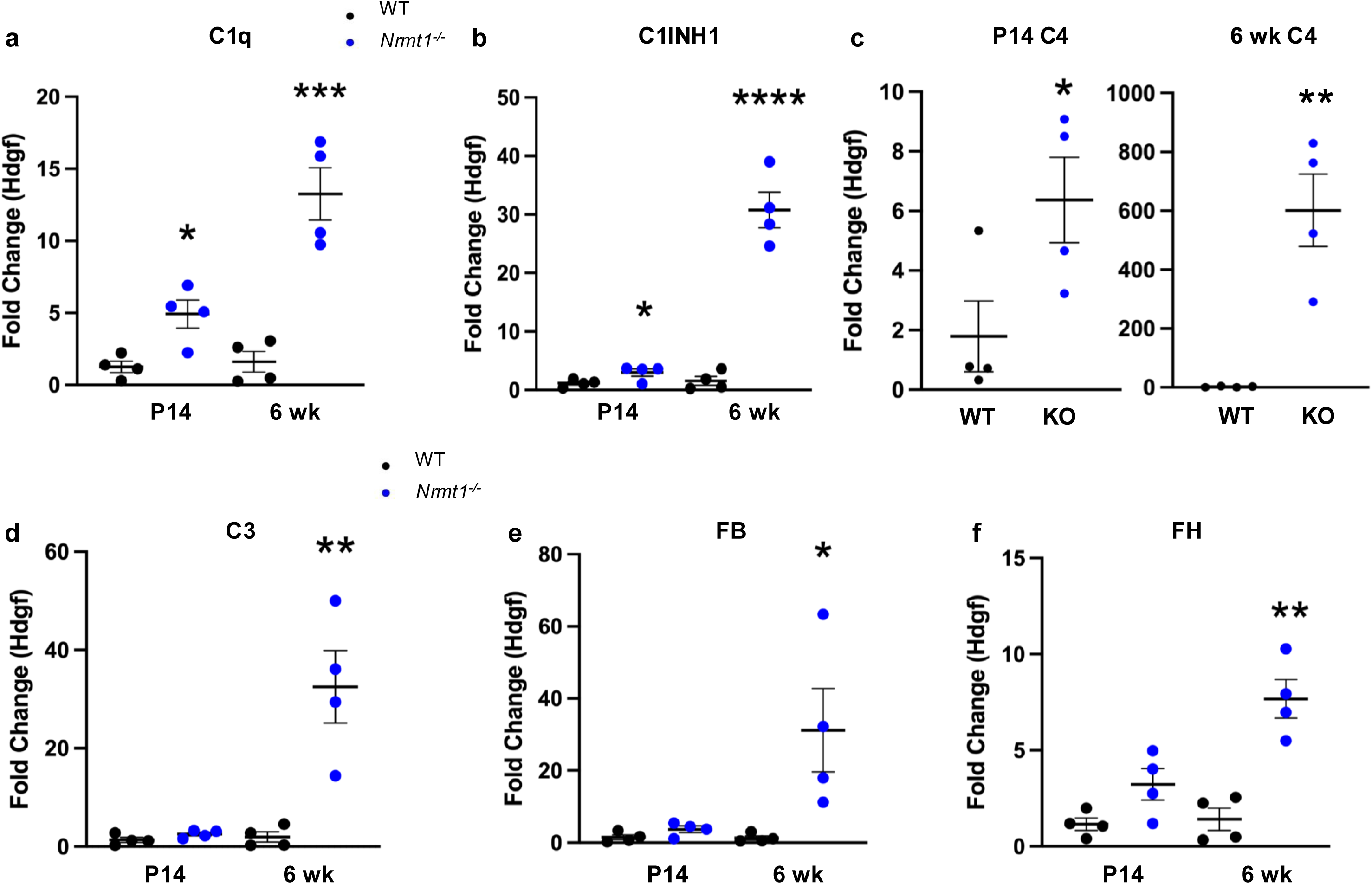
qRT-PCR analysis of Complement protein expression. **a-c** *Nrmt1^-/-^* mice exhibit significant upregulation of transcripts involved in classical pathway of complement cascade beginning at P14 and increasing at 6 weeks. **d-f** Transcripts involved in the alternative pathway of the complement cascade do not become significantly upregulated until 6 weeks in *Nrmt1^-/-^* mice. *p<0.05, **p<0.01, ***p<0.001, ****p<0.0001 as determined by unpaired t-test and error bars signify mean±SEM. n=4.

### Blood brain barrier and anti-inflammatory response

As activation of C3 can result in decreased blood brain barrier (BBB) integrity and increased infiltration of inflammatory cells into the brain (56), we also used qPCR to assay for markers of the BBB integrity. During neuroinflammation the permeability of the BBB increases to allow for the infiltration of peripheral immune cells, which can release both pro-and anti-inflammatory cytokines, capable of exacerbating and resolving neuroinflammatory responses (44, 57–60). To determine if BBB integrity may be disrupted in *Nrmt1^-/-^* mice, we used qRT-PCR in striatal lysates from P14 and 6-week *Nrmt1^-/-^* and WT mice to assess the mRNA expression of the extracellular matrix (ECM) reorganization proteins, tissue inhibitors of metalloproteinase-1 (TIMP1) and matrix metalloproteinase-9 (MMP9), as well as the tight-junctional protein claudin-5. We observed a significant increase in *Timp1* mRNA expression beginning at P14 and persisting, becoming increasingly more significant at 6 weeks (**Fig. 6a**). However, mRNA expression of *Mmp9* and *claudin-5* was not significantly upregulated until 6-weeks (**Fig. 6b,c).** Increases in *Timp1* and *Mmp9* expression are associated with changes in ECM homeostasis, BBB leakage, and neurodegenerative diseases (61–63). Though increases in *claudin-5* expression can signify decreased BBB permeability and leakage of proteins from circulation (57, 62), the observed increase in its expression at 6-weeks could also suggest a potential compensatory mechanism in effort to maintain BBB integrity.

**Fig. 6.**
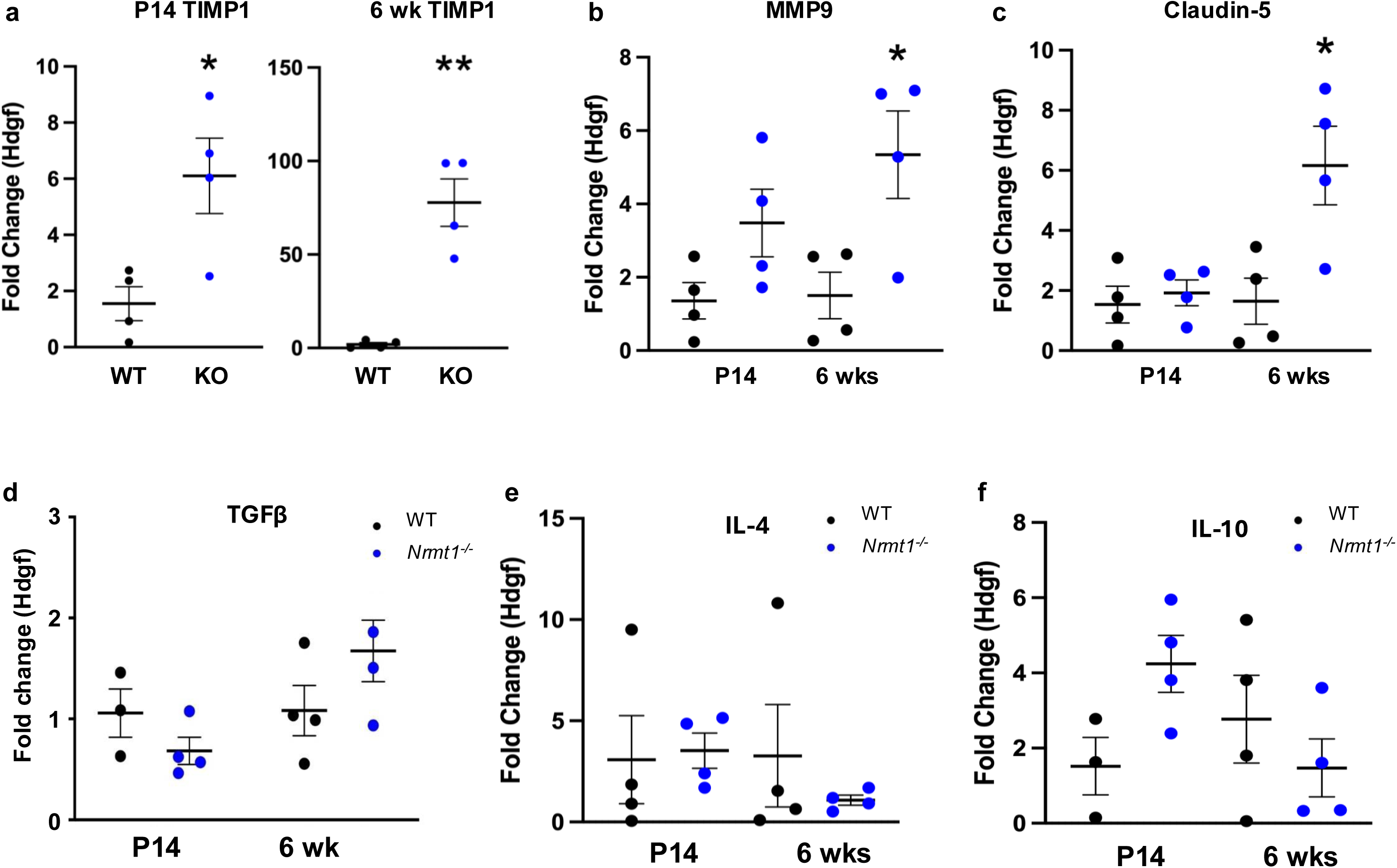
qPCR analysis of BBB integrity and anti-inflammatory markers. **a** *Timp1* expression is significantly increased in the striatum of *Nrmt1^-/-^*mice at P14 and 6 weeks as compared to wild type (WT) controls. Though **b**) *Mmp9* and **c**) *claudin-5* expression are only significantly increased at 6 weeks. **d-f** No differences in expression were seen in the anti-inflammatory markers *Tgfβ, Il-4,* or *Il-10*. *p<0.05, **p<0.01 as determined by unpaired t-test and error bars signify mean±SEM. n=4.

Finally, resolution of a neuroinflammatory response is critical in preventing chronic inflammation and tissue damage, so pro-inflammatory cytokine release often triggers the release of counteracting anti-inflammatory cytokines, and dysregulation of this balance has been proposed to contribute neurodegenerative phenotypes (44, 64, 65). Anti-inflammatory cytokines, such as TGFβ, IL-4 and IL-10, play a major role in balancing immune responses and are capable of preventing chronic inflammation, microglia activation, and neuron loss in animal models (66–73). To determine if *Nrmt1^-/-^* mice can mount an anti-inflammatory response, we used qRT-PCR to assess the mRNA expression of the anti-inflammatory cytokines TGFβ, IL-4, and IL-10 in striatal lysates at P14 and 6 weeks. We observed no significant differences in the expression of *Tgfβ*, *Il-4*, and *Il-10* in the striatum of *Nrmt1^-/-^* mice compared to WT controls at either age (**Fig. 6d-f**), indicating that despite the significant pro-inflammatory response observed in *Nrmt1^-/-^* mice, a counteracting anti-inflammatory response is not activated. Together, these results suggest that, though *Nrmt1^-/-^*mice may have increased infiltration of peripheral immune cells, they cannot activate an anti-inflammatory response, leading to an inability to properly balance inflammatory reactions.

## Discussion

Our findings reveal a clear mechanistic link between early neurogenic defects and subsequent neurodegeneration in *Nrmt1^⁻/⁻^* mice, mediated by sustained neuroinflammation. Aberrant neural stem cell cycle regulation is followed by p25/CDK5 hyperactivation, initiation of pro-inflammatory cytokine signaling, complement pathway activation, microgliosis, astrogliosis, and potential blood-brain barrier (BBB) irregularities, highlighting how a developmental abnormality can set in motion a self-reinforcing degenerative cascade driven by inflammation. That *Nrmt1^⁻/⁻^* mice may not be able to mount a counteracting anti-inflammatory response, could explain why progressive neurodegeneration is observed at a relatively young age in this model.

The early persistence of actively cycling NeuN⁺ neurons at P14 underscores a fundamental defect in postnatal neuronal maturation (29). Incomplete cell cycle exit is a known trigger for neuronal vulnerability, as inappropriate re-entry into the cycle can activate apoptotic pathways (30). The subsequent rise in p25 levels by 6 weeks, absent in WT controls, indicates that neurotoxic CDK5 hyperactivation is a downstream consequence of these developmental errors. Tau hyperphosphorylation at Ser202, an established substrate of p25/CDK5, further implicates this pathway in cytoskeletal destabilization and synaptic dysfunction (39), hallmarks of many neurodegenerative disorders.

The inflammatory profile observed here mirrors the trajectory seen in human disease, early TNF upregulation followed by later increases in IL-1 and IL-6, marking the shift from initial immune activation to a chronic pro-inflammatory state (64, 74). The emergence of astrogliosis between 4 and 6 weeks, closely paralleling neuronal loss, suggests that reactive astrocytes may act as both responders to injury and amplifiers of inflammation, possibly via the TNF/STAT3 axis (46, 60). The lack of compensatory anti-inflammatory cytokines such as IL-10 or TGFβ indicates that the homeostatic M2 microglial phenotype fails to counterbalance the dominant M1 state, thereby permitting sustained damage (68, 73). Complement activation emerged as an especially early event, with classical pathway components (C1q, C1INH, C4) upregulated at P14, well before widespread neurodegeneration. Interestingly, mRNA expression of complement C3 is not upregulated in the striatum at P14, suggesting the increase in C3 immunoreactivity may be due to post-transcriptional regulation or increased protein stability and accumulation, rather than transcriptional upregulation at this time point. This finding highlights a potentially interesting aspect of C3 regulation in *Nrmt1^⁻/⁻^* mice that warrants further investigation. The detection of C3 deposition by 4 weeks, coinciding with a marker of microgliosis, suggests that complement-mediated synaptic tagging may prime neural circuits for microglial engulfment and loss, as documented in Alzheimer’s and Huntington’s disease models (49, 50). This temporal precedence raises the possibility that complement activation is a driver, rather than merely a marker, of degeneration in *Nrmt1^⁻/⁻^* mice(75, 76).

The altered expression of BBB remodeling markers further supports a transition toward a permissive environment for peripheral immune infiltration. Early TIMP1 upregulation followed by MMP9 elevation and increased claudin-5 expression suggests a dynamic, but ultimately insufficient, attempt to preserve barrier integrity (57, 77). These changes are likely to exacerbate neuronal vulnerability by allowing pro-inflammatory mediators and immune cells into the CNS, further amplifying injury (58, 61). The lack of an anti-inflammatory response under these conditions suggests that *Nrmt1^⁻/⁻^* mice may also have hematopoietic differentiation defects that compound the differentiation defects in other tissues.

The *Nrmt1^⁻/⁻^* mouse provides a unique model for studying how impaired neurogenesis initiates neuroinflammatory processes that culminate in neuronal loss. As this neuronal loss happens early in life, these mice can also be used to identify missing compensatory mechanisms that mask symptoms until later in life. By bridging the gap between developmental biology and neurodegeneration, these findings underscore the need to identify and target early vulnerabilities such as aberrant neuronal maturation or excessive complement activation before they cascade into irreversible neurodegeneration. Therapeutic strategies that restore cell cycle fidelity, modulate complement activity, or rebalance microglial phenotypes toward anti-inflammatory states may hold promise for delaying or preventing late-onset neurodegenerative disease.

## Acknowledgements

This work was supported by a research grant from the National Institutes of Health to CST [GM144111].

---

**Sup. Fig. 1.**
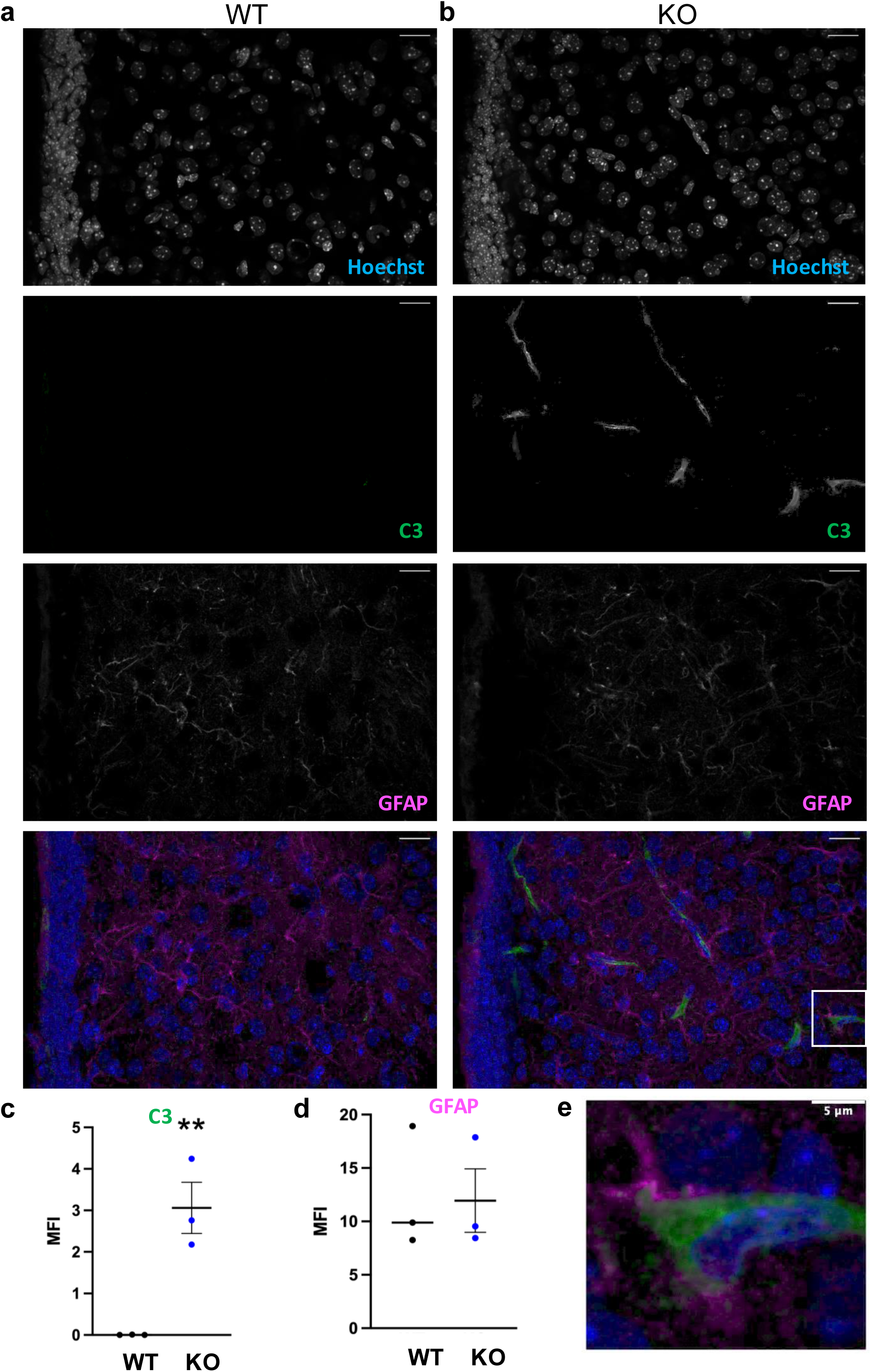
Co-localization of C3 and GFAP staining at P14. **a** Minimal C3 staining is seen in wild type (WT) mice at P14. **b** The diffuse C3 staining seen in P14 *Nrmt1^-/-^* mice overlaps with the processes of GFAP+ cells. **c,d** Quantification of C3 and GFAP staining. **e** Enlargement of cell in white box above. Scale bars represent 20 μm unless otherwise stated. **p<0.01 as determined by unpaired t-test and error bars signify mean±SEM.

## References

1. Hou Y, Dan X, Babbar M, Wei Y, Hasselbalch SG, Croteau DL, et al. Ageing as a risk factor for neurodegenerative disease. Nature reviews Neurology. 2019;15(10):565–81.

2. Galgoczi S, Ruzo A, Markopoulos C, Yoney A, Phan-Everson T, Li S, et al. Huntingtin CAG expansion impairs germ layer patterning in synthetic human 2D gastruloids through polarity defects. Development (Cambridge, England). 2021;148(19).

3. Capizzi M, Carpentier R, Denarier E, Adrait A, Kassem R, Mapelli M, et al. Developmental defects in Huntington’s disease show that axonal growth and microtubule reorganization require NUMA1. Neuron. 2022;110(1):36–50.e5.

4. Barnat M, Capizzi M, Aparicio E, Boluda S, Wennagel D, Kacher R, et al. Huntington’s disease alters human neurodevelopment. Science (New York, NY). 2020;369(6505):787-93.

5. Wiatr K, Szlachcic WJ, Trzeciak M, Figlerowicz M, Figiel M. Huntington Disease as a Neurodevelopmental Disorder and Early Signs of the Disease in Stem Cells. Molecular neurobiology. 2018;55(4):3351–71.

6. Töpper R, Schwarz M, Lange HW, Hefter H, Noth J. Neurophysiological abnormalities in the Westphal variant of Huntington’s disease. Movement disorders: official journal of the Movement Disorder Society. 1998;13(6):920–8.

7. Ruzo A, Croft GF, Metzger JJ, Galgoczi S, Gerber LJ, Pellegrini C, et al. Chromosomal instability during neurogenesis in Huntington’s disease. Development (Cambridge, England). 2018;145(2).

8. Garcia-Reitboeck P, Anichtchik O, Dalley JW, Ninkina N, Tofaris GK, Buchman VL, et al. Endogenous alpha-synuclein influences the number of dopaminergic neurons in mouse substantia nigra. Experimental neurology. 2013;248:541–5.

9. Wang J, Gallagher D, DeVito LM, Cancino GI, Tsui D, He L, et al. Metformin activates an atypical PKC-CBP pathway to promote neurogenesis and enhance spatial memory formation. Cell stem cell. 2012;11(1):23–35.

10. Anichtchik O, Diekmann H, Fleming A, Roach A, Goldsmith P, Rubinsztein DC. Loss of PINK1 function affects development and results in neurodegeneration in zebrafish. The Journal of neuroscience: the official journal of the Society for Neuroscience. 2008;28(33):8199–207.

11. Bahnassawy L, Nicklas S, Palm T, Menzl I, Birzele F, Gillardon F, et al. The parkinson’s disease-associated LRRK2 mutation R1441G inhibits neuronal differentiation of neural stem cells. Stem cells and development. 2013;22(18):2487–96.

12. Scarmeas N, Habeck CG, Stern Y, Anderson KE. APOE genotype and cerebral blood flow in healthy young individuals. Jama. 2003;290(12):1581–2.

13. Freude KK, Penjwini M, Davis JL, LaFerla FM, Blurton-Jones M. Soluble amyloid precursor protein induces rapid neural differentiation of human embryonic stem cells. The Journal of biological chemistry. 2011;286(27):24264–74.

14. Sapir T, Frotscher M, Levy T, Mandelkow EM, Reiner O. Tau’s role in the developing brain: implications for intellectual disability. Hum Mol Genet. 2012;21(8):1681–92.

15. Etoc F, Metzger J, Ruzo A, Kirst C, Yoney A, Ozair MZ, et al. A Balance between Secreted Inhibitors and Edge Sensing Controls Gastruloid Self-Organization. Dev Cell. 2016;39(3):302–15.

16. Marchetti B. Wnt/β-Catenin Signaling Pathway Governs a Full Program for Dopaminergic Neuron Survival, Neurorescue and Regeneration in the MPTP Mouse Model of Parkinson’s Disease. International journal of molecular sciences. 2018;19(12).

17. Beatus P, Lendahl U. Notch and neurogenesis. J Neurosci Res. 1998;54(2):125–36.

18. van der Plas E, Schultz JL, Nopoulos PC. The Neurodevelopmental Hypothesis of Huntington’s Disease. J Huntingtons Dis. 2020;9(3):217–29.

19. Bayly RD, Brown CY, Agarwala S. A novel role for FOXA2 and SHH in organizing midbrain signaling centers. Developmental biology. 2012;369(1):32–42.

20. Distéfano-Gagné F, Bitarafan S, Lacroix S, Gosselin D. Roles and regulation of microglia activity in multiple sclerosis: insights from animal models. Nature reviews Neuroscience. 2023;24(7):397–415.

21. Hansen DV, Hanson JE, Sheng M. Microglia in Alzheimer’s disease. The Journal of cell biology. 2018;217(2):459–72.

22. Yeh FL, Wang Y, Tom I, Gonzalez LC, Sheng M. TREM2 Binds to Apolipoproteins, Including APOE and CLU/APOJ, and Thereby Facilitates Uptake of Amyloid-Beta by Microglia. Neuron. 2016;91(2):328–40.

23. Samuels JD, Lukens JR, Price RJ. Emerging roles for ITAM and ITIM receptor signaling in microglial biology and Alzheimer’s disease-related amyloidosis. Journal of neurochemistry. 2024;168(10):3558–73.

24. Song GJ, Suk K. Pharmacological Modulation of Functional Phenotypes of Microglia in Neurodegenerative Diseases. Frontiers in aging neuroscience. 2017;9:139.

25. Cui W, Sun C, Ma Y, Wang S, Wang X, Zhang Y. Inhibition of TLR4 Induces M2 Microglial Polarization and Provides Neuroprotection via the NLRP3 Inflammasome in Alzheimer’s Disease. Frontiers in neuroscience. 2020;14:444.

26. Fonseca MI, Chu SH, Hernandez MX, Fang MJ, Modarresi L, Selvan P, et al. Cell-specific deletion of C1qa identifies microglia as the dominant source of C1q in mouse brain. J Neuroinflammation. 2017;14(1):48.

27. Schafer DP, Lehrman EK, Kautzman AG, Koyama R, Mardinly AR, Yamasaki R, et al. Microglia sculpt postnatal neural circuits in an activity and complement-dependent manner. Neuron. 2012;74(4):691–705.

28. Bonsignore LA, Tooley JG, Van Hoose PM, Wang E, Cheng A, Cole MP, et al. NRMT1 knockout mice exhibit phenotypes associated with impaired DNA repair and premature aging. Mechanisms of ageing and development. 2015;146–148:42-52.

29. Catlin JP, Marziali LN, Rein B, Yan Z, Feltri ML, Schaner Tooley CE. Age-related neurodegeneration and cognitive impairments of NRMT1 knockout mice are preceded by misregulation of RB and abnormal neural stem cell development. Cell death & disease. 2021;12(11):1014.

30. Barrio-Alonso E, Hernández-Vivanco A, Walton CC, Perea G, Frade JM. Cell cycle reentry triggers hyperploidization and synaptic dysfunction followed by delayed cell death in differentiated cortical neurons. Scientific reports. 2018;8(1):14316.

31. Vacher H, Yang JW, Cerda O, Autillo-Touati A, Dargent B, Trimmer JS. Cdk-mediated phosphorylation of the Kvβ2 auxiliary subunit regulates Kv1 channel axonal targeting. The Journal of cell biology. 2011;192(5):813–24.

32. Lee MS, Kwon YT, Li M, Peng J, Friedlander RM, Tsai LH. Neurotoxicity induces cleavage of p35 to p25 by calpain. Nature. 2000;405(6784):360-4.

33. Maitra S, Vincent B. Cdk5-p25 as a key element linking amyloid and tau pathologies in Alzheimer’s disease: Mechanisms and possible therapeutic interventions. Life Sci. 2022;308:120986.

34. Tran J, Taylor SKB, Gupta A, Amin N, Pant H, Gupta BP, et al. Therapeutic effects of TP5, a Cdk5/p25 inhibitor, in in vitro and in vivo models of Parkinson’s disease. Curr Res Neurobiol. 2021;2:100006.

35. Wilkaniec A, Czapski GA, Adamczyk A. Cdk5 at crossroads of protein oligomerization in neurodegenerative diseases: facts and hypotheses. Journal of neurochemistry. 2016;136(2):222–33.

36. Paoletti P, Vila I, Rifé M, Lizcano JM, Alberch J, Ginés S. Dopaminergic and glutamatergic signaling crosstalk in Huntington’s disease neurodegeneration: the role of p25/cyclin-dependent kinase 5. The Journal of neuroscience: the official journal of the Society for Neuroscience. 2008;28(40):10090–101.

37. Tripathi RB, Jackiewicz M, McKenzie IA, Kougioumtzidou E, Grist M, Richardson WD. Remarkable Stability of Myelinating Oligodendrocytes in Mice. Cell reports. 2017;21(2):316–23.

38. Lehotzky A, Lau P, Tokési N, Muja N, Hudson LD, Ovádi J. Tubulin polymerization-promoting protein (TPPP/p25) is critical for oligodendrocyte differentiation. Glia. 2010;58(2):157–68.

39. Hashiguchi M, Saito T, Hisanaga S, Hashiguchi T. Truncation of CDK5 activator p35 induces intensive phosphorylation of Ser202/Thr205 of human tau. The Journal of biological chemistry. 2002;277(46):44525–30.

40. Sharma A, Brenner M, Jacob A, Marambaud P, Wang P. Extracellular CIRP Activates the IL-6Rα/STAT3/Cdk5 Pathway in Neurons. Molecular neurobiology. 2021;58(8):3628–40.

41. Sundaram JR, Chan ES, Poore CP, Pareek TK, Cheong WF, Shui G, et al. Cdk5/p25-induced cytosolic PLA2-mediated lysophosphatidylcholine production regulates neuroinflammation and triggers neurodegeneration. The Journal of neuroscience: the official journal of the Society for Neuroscience. 2012;32(3):1020–34.

42. Parameswaran N, Patial S. Tumor necrosis factor-α signaling in macrophages. Crit Rev Eukaryot Gene Expr. 2010;20(2):87–103.

43. Cruz JC, Tseng HC, Goldman JA, Shih H, Tsai LH. Aberrant Cdk5 activation by p25 triggers pathological events leading to neurodegeneration and neurofibrillary tangles. Neuron. 2003;40(3):471–83.

44. Kempuraj D, Thangavel R, Natteru PA, Selvakumar GP, Saeed D, Zahoor H, et al. Neuroinflammation Induces Neurodegeneration. J Neurol Neurosurg Spine. 2016;1(1).

45. Pekna M, Pekny M. The Complement System: A Powerful Modulator and Effector of Astrocyte Function in the Healthy and Diseased Central Nervous System. Cells. 2021;10(7).

46. Lawrence JM, Schardien K, Wigdahl B, Nonnemacher MR. Roles of neuropathology-associated reactive astrocytes: a systematic review. Acta neuropathologica communications. 2023;11(1):42.

47. Chen Y, Chu JMT, Chang RCC, Wong GTC. The Complement System in the Central Nervous System: From Neurodevelopment to Neurodegeneration. Biomolecules. 2022;12(2):337.

48. Sakai J. Core Concept: How synaptic pruning shapes neural wiring during development and, possibly, in disease. Proc Natl Acad Sci U S A. 2020;117(28):16096–9.

49. Hong S, Beja-Glasser VF, Nfonoyim BM, Frouin A, Li S, Ramakrishnan S, et al. Complement and microglia mediate early synapse loss in Alzheimer mouse models. Science (New York, NY). 2016;352(6286):712-6.

50. Wilton DK, Mastro K, Heller MD, Gergits FW, Willing CR, Fahey JB, et al. Microglia and complement mediate early corticostriatal synapse loss and cognitive dysfunction in Huntington’s disease. Nat Med. 2023;29(11):2866–84.

51. Butler CA, Popescu AS, Kitchener EJA, Allendorf DH, Puigdellívol M, Brown GC. Microglial phagocytosis of neurons in neurodegeneration, and its regulation. J Neurochem. 2021;158(3):621–39.

52. Dejanovic B, Wu T, Tsai MC, Graykowski D, Gandham VD, Rose CM, et al. Complement C1q-dependent excitatory and inhibitory synapse elimination by astrocytes and microglia in Alzheimer’s disease mouse models. Nat Aging. 2022;2(9):837–50.

53. Clark DPQ, Perreau VM, Shultz SR, Brady RD, Lei E, Dixit S, et al. Inflammation in Traumatic Brain Injury: Roles for Toxic A1 Astrocytes and Microglial-Astrocytic Crosstalk. Neurochem Res. 2019;44(6):1410–24.

54. Zhou R, Chen SH, Zhao Z, Tu D, Song S, Wang Y, et al. Complement C3 Enhances LPS-Elicited Neuroinflammation and Neurodegeneration Via the Mac1/NOX2 Pathway. Molecular neurobiology. 2023;60(9):5167–83.

55. Zemtsova I, Görg B, Keitel V, Bidmon HJ, Schrör K, Häussinger D. Microglia activation in hepatic encephalopathy in rats and humans. Hepatology (Baltimore, Md). 2011;54(1):204–15.

56. Wu F, Zou Q, Ding X, Shi D, Zhu X, Hu W, et al. Complement component C3a plays a critical role in endothelial activation and leukocyte recruitment into the brain. J Neuroinflammation. 2016;13:23.

57. Kadry H, Noorani B, Cucullo L. A blood-brain barrier overview on structure, function, impairment, and biomarkers of integrity. Fluids Barriers CNS. 2020;17(1):69.

58. Knox EG, Aburto MR, Clarke G, Cryan JF, O’Driscoll CM. The blood-brain barrier in aging and neurodegeneration. Molecular psychiatry. 2022;27(6):2659–73.

59. Propson NE, Roy ER, Litvinchuk A, Köhl J, Zheng H. Endothelial C3a receptor mediates vascular inflammation and blood-brain barrier permeability during aging. Journal of Clinical Investigation. 2021;131(1).

60. Kim H, Leng K, Park J, Sorets AG, Kim S, Shostak A, et al. Reactive astrocytes transduce inflammation in a blood-brain barrier model through a TNF-STAT3 signaling axis and secretion of alpha 1-antichymotrypsin. Nature communications. 2022;13(1):6581.

61. Taylor X, Clark IM, Fitzgerald GJ, Oluoch H, Hole JT, DeMattos RB, et al. Amyloid-β (Aβ) immunotherapy induced microhemorrhages are associated with activated perivascular macrophages and peripheral monocyte recruitment in Alzheimer’s disease mice. Molecular neurodegeneration. 2023;18(1):59.

62. Rempe RG, Hartz AMS, Bauer B. Matrix metalloproteinases in the brain and blood-brain barrier: Versatile breakers and makers. J Cereb Blood Flow Metab. 2016;36(9):1481–507.

63. Ashutosh, Chao C, Borgmann K, Brew K, Ghorpade A. Tissue inhibitor of metalloproteinases-1 protects human neurons from staurosporine and HIV-1-induced apoptosis: mechanisms and relevance to HIV-1-associated dementia. Cell Death & Disease. 2012;3(6):e332-e.

64. Heneka MT, Carson MJ, El Khoury J, Landreth GE, Brosseron F, Feinstein DL, et al. Neuroinflammation in Alzheimer’s disease. Lancet Neurol. 2015;14(4):388–405.

65. Schwartz M, Deczkowska A. Neurological Disease as a Failure of Brain-Immune Crosstalk: The Multiple Faces of Neuroinflammation. Trends Immunol. 2016;37(10):668–79.

66. Bido S, Nannoni M, Muggeo S, Gambarè D, Ruffini G, Bellini E, et al. Microglia-specific IL-10 gene delivery inhibits neuroinflammation and neurodegeneration in a mouse model of Parkinson’s disease. Sci Transl Med. 2024;16(761):eadm8563.

67. Gadani SP, Cronk JC, Norris GT, Kipnis J. IL-4 in the brain: a cytokine to remember. J Immunol. 2012;189(9):4213–9.

68. Gao Y, Tu D, Yang R, Chu CH, Hong JS, Gao HM. Through Reducing ROS Production, IL-10 Suppresses Caspase-1-Dependent IL-1β Maturation, thereby Preventing Chronic Neuroinflammation and Neurodegeneration. International journal of molecular sciences. 2020;21(2).

69. Norden DM, Fenn AM, Dugan A, Godbout JP. TGFβ produced by IL-10 redirected astrocytes attenuates microglial activation. Glia. 2014;62(6):881–95.

70. Porro C, Cianciulli A, Panaro MA. The Regulatory Role of IL-10 in Neurodegenerative Diseases. Biomolecules. 2020;10(7).

71. Russo K, Wharton KA. BMP/TGF-β signaling as a modulator of neurodegeneration in ALS. Dev Dyn. 2022;251(1):10–25.

72. Xie L, Liu Y, Zhang N, Li C, Sandhu AF, Williams G, 3rd, et al. Electroacupuncture Improves M2 Microglia Polarization and Glia Anti-inflammation of Hippocampus in Alzheimer’s Disease. Front Neurosci. 2021;15:689629.

73. Lobo-Silva D, Carriche GM, Castro AG, Roque S, Saraiva M. Balancing the immune response in the brain: IL-10 and its regulation. J Neuroinflammation. 2016;13(1):297.

74. Hampel H, Caraci F, Cuello AC, Caruso G, Nisticò R, Corbo M, et al. A Path Toward Precision Medicine for Neuroinflammatory Mechanisms in Alzheimer’s Disease. Front Immunol. 2020;11:456.

75. Tenner AJ, Stevens B, Woodruff TM. New tricks for an ancient system: Physiological and pathological roles of complement in the CNS. Mol Immunol. 2018;102:3–13.

76. Stephan AH, Barres BA, Stevens B. The complement system: an unexpected role in synaptic pruning during development and disease. Annual review of neuroscience. 2012;35:369–89.

77. Tang J, Kang Y, Huang L, Wu L, Peng Y. TIMP1 preserves the blood-brain barrier through interacting with CD63/integrin β 1 complex and regulating downstream FAK/RhoA signaling. Acta Pharm Sin B. 2020;10(6):987–1003.

